# Identifiability of pharmacological models for online individualization

**DOI:** 10.1101/2021.11.03.467092

**Authors:** Ylva Wahlquist, Amina Gojak, Kristian Soltesz

## Abstract

There is a large variability between individuals in the response to anesthetic drugs, that seriously limits the achievable performance of closed-loop controlled drug dosing. Full individualization of patient models based on early clinical response data has been suggested as a means to improve performance with maintained robustness (safety). We use estimation theoretic analysis and realization theory to characterize practical identifiability of the standard pharmacological model structure from anesthetic induction phase data and conclude that such approaches are not practically feasible.

## 1. INTRODUCTION

In general anesthesia, hypnotic depth—reciprocal to the level of awareness—is controlled through the addition of anesthetic drugs distributed across tissues and eventually metabolized. This paper considers the prospect of automatically controlling the hypnotic depth through real-time electroencephalogram (EEG) measurements, from which the depth can be estimated, and intravenous infusion of the anesthetic drug propofol. This setting is well-studied in the literature and has been the subject of a handful of clinical studies. Without attempting a comprehensive survey, we suggest the survey Ghita et al. (2000) as a starting point for the interested reader.

As with many drugs, there is a large inter-patient variability in the response to propofol. It can partly be explained using mixed-effect population models as described in Eleveld et al. (2018). Even so, the remaining variability remains large, and importantly limits the performance of a safe (robust) feedback controller, as we have illustrated in for example Gonzales-Cava et al. (2020).

Consequently, several simulation studies have suggested to individualize the controller, based on data from the induction phase of anesthesia, comprising of the transition from the fully aware state to a desired hypnotic depth. In Soltesz et al. (2013) we obtained promising results when considering an idealized simulation case. Here, we return to the same idea and investigate how including a measurement noise model changes the prospect of individualized therapy based induction phase data. Particularly, we investigate the practical identifiability of the standard pharmacological model used to describe the effect of propofol on the hypnotic depth from induction phase time series. This is what would be required in a clinical setting since the point of the individualization is to have the model available for updating the drug dosing controller as early as possible during the ongoing treatment.

## 2. DYNAMICAL MODEL

### 2.1. Pharmacological model

The relation between propofol infusion *u* [mass/time] and measured clinical effect *y* is commonly modeled by a pharmacokinetic (PK) model relating *u* to the blood plasma concentration *C*_*p*_ [mass/volume], a pharmacodynamic (PD) model relating *C*_*p*_ to the (normalized) clinical effect 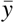, and finally an observation model relating 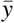 to the measured clinical effect *y*.

The combined PK and PD (PKPD) model comprises a linear and time invariant (LTI) mammillary three-compartment model with *C*_*p*_ as output, connected in series with a lowpass filter, that can be thought of as an effect site compartment of infinitesimal volume. The output of the series connection is the effect site concentration *C*_*e*_ [mass/volume]. Here we will normalize *C*_*e*_ by *EC*_50_ [mass/volume], being the *C*_*e*_ value corresponding to the midpoint between no and full clinical effect. We thus introduce *z* = *C*_*e*_ */EC*_50_. The combined LTI parts of the PKPD models can be expressed on state space form as

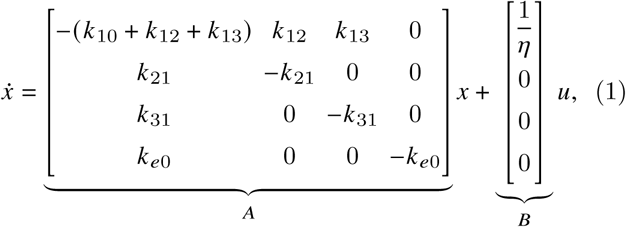

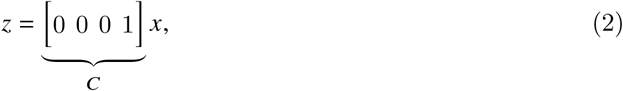

where *η* = *v*_1_*EC*_50_ is the product of the central compartment volume *v*_1_ [volume] and *EC*_50_.

The transfer function of (1)–(2) from *u* to *z* is

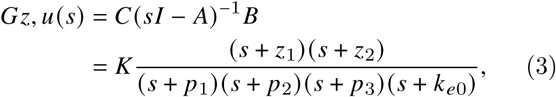

with static gain *K* = *k*_*e*0_ /*η*, zeros − *z*_1_ = − *k*_21_, − *z*_2_ = − *k*_31_, and where the poles − *p*_1_, − *p*_2_, − *p*_3_ solve the characteristic equations

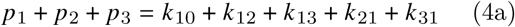

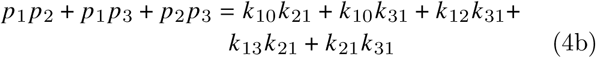

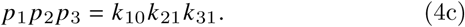

The nonlinear output part of the PD model is characterized by a sigmoidal function *h*, relating *z* = *C*_*e*_ / *EC*_50_ to the normalized clinical effect:

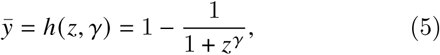

where 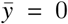 and 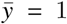 signify full awareness, and the maximal hypnotic depth, respectively.

The eight parameters *k*_10_, *k*_12_, *k*_13_, *k*_21_, *k*_31_, *k*_*e*0_, *γ, η* are assumed strictly positive, with *γ* ≥ 1. The six rate parameters *k*: [1/time] define the diffusion between the compartments; *η* has unit [mass]; *γ* is unitless and defines the shape of (5). A more in-depth explanation of the combined model (1)–(5) is provided in Copot (2019).

### 2.2. Actuation model

The input *u* is actuated using a computer-controlled infusion pump. For any reasonable syringe drug concentration quantization error in the magnitude of *u(t)* is negligible, as is quantization in *t* introduced by zero-order-hold (ZOH) sampling ^1^ Furthermore, since the input is ZOH, ZOH sampling of (1) at *T*_*s*_ is approximation-free and used for all simulations herein.

### 2.3. Observation model

Clinicians are used to the awareness being reported using the bispectral index (BIS) or other awareness estimates reported on the same scale ^2^, where

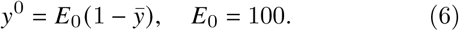

We will here consider *E*_0_ = 100 in the equilibrium state *x* = 0, but it could alternatively be considered a free parameter that accounts for some patients generating a monitor reading of e.g. 90 BIS (by setting *E*_0_ = 90) in absence of drug, as observed and modeled in van Heusden et al. (2013).

The clinical effect measure *y* is an *estimate* of the clinical effect *y*^0^, derived from EEG measurements using a monitoring device, with a ZOH output, updated at *T*_*s*_ = 1 s. The observation model employed here is that the monitor introduces additive independent and identically distributed (IID) Gaussian noise *w* ∼ 𝒩 (0, *σ*^2^),

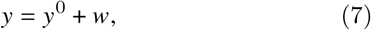

where *σ*^2^ = 9^2^ BIS was identified for the NeuroSense EEG monitor in Soltesz et al. (2012). Herein we use the more optimistic *σ*^2^ = 1 BIS ^3^.

## 3. SIMULATED EXAMPLE

To generate a representative virtual patient, we have used the population model of Eleveld et al. (2018) that, based on openly available data sets, defines (nominal) individual models on the form (1)–(5) as functions of patient demographics. Particularly for our investigation, we have used the reference patient model (35 year old male, 1.70 m, 70 kg) with parameter values *θ*° in table 1.

**Table 1.** Parameters (left) together with time constants 1/*p*: and 1/*z*: (right) of the system poles and zeros of the model (1)–(5) corresponding to Fig. 2–3.

| | $k_{10}$ | $k_{12}$ | $k_{13}$ | $k_{21}$ | $k_{31}$ | $k_{e0}$ | $\gamma$ | $\eta$ | $1/p_1$ | $1/p_2$ | $1/p_3$ | $1/k_{e0}$ | $1/z_1$ | $1/z_2$ |
| --- | --- | --- | --- | --- | --- | --- | --- | --- | --- | --- | --- | --- | --- | --- |
| Unit | $10^3/\text{s}$ | | | | | | 1 | $\mu\text{g}$ | min | | | | | |
| $\theta^o$ | 4.8 | 8.6 | 2.9 | 2.1 | 0.068 | 2.4 | 1.47 | 19342 | 0.95 | 18 | 406 | 6.8 | 7.9 | 246 |
| $\Delta k_{10}$ | 475 | 7626 | 243 | 213 | 0.72 | 1.3 | 1.66 | 267 | 0.0020 | 0.92 | 35 | 13 | 0.078 | 23 |
| $\Delta k_{12}$ | 0.94 | 0.086 | 0.17 | 2.9 | 0.54 | 18 | 1.48 | $1.4\text{e}5$ | 5.5 | 14 | 41 | 0.93 | 5.7 | 31 |
| $\Delta k_{13}$ | 12 | 5.6 | 0.029 | 0.79 | 0.79 | 1.3 | 1.48 | 10857 | 0.92 | 31 | 96 | 12 | 21 | 96 |
| $\Delta k_{21}$ | 0.95 | $9.2\text{e-}10$ | 0.19 | 212 | 0.79 | 18 | 1.48 | $1.4\text{e}5$ | 0.079 | 12 | 31 | 0.92 | 0.08 | 21 |
| $\Delta k_{31}$ | 1.0 | 0.18 | 0.63 | 0.47 | 6.8 | 16 | 1.45 | $1.3\text{e}5$ | 2.2 | 14 | 46 | 1.0 | 2.5 | 35 |
| $\Delta k_{e0}$ | $1.5\text{e-}11$ | $1.2\text{e-}8$ | 0.53 | 2647 | 0.030 | 243 | 5.1 | $2.0\text{e}5$ | 0.0063 | 30 | $2.1\text{e}13$ | 0.069 | 0.0063 | 558 |
| $\Delta\gamma$ | 17 | 1222 | 0.25 | 26 | 0.49 | 25 | 147 | 4733 | 0.014 | 32 | 47 | 0.66 | 0.65 | 34 |
| $\Delta\eta$ | 687 | 60600 | 316 | 1124 | 0.77 | 1.4 | 1.51 | 193 | $3.0\text{e-}4$ | 0.91 | 32 | 12 | 0.015 | 22 |

The experiment is commenced in the stationary state *x* = 0 of (1), corresponding to the drug-free equilibrium. At time *t* = 0 a bolus dose (an impulse) *αδ*(0) is delivered and a steady infusion *β* is commenced. The input is thus

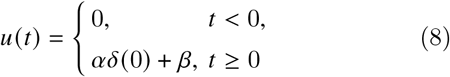

The steady infusion was chosen to yield 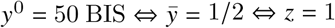 in stationarity. With system matrices defined through (1)–(2) this corresponds to solving −*CA*^−1^*Bβ* = 1, resulting in *β* = *k*_10_ *η* = 5.5 mg/min.

In our simulations, the bolus was approximated by a high-rate infusion ^4^ and the time during which the bolus dose was delivered was chosen (through bisection search) to limit the response overshoot to 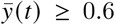 as motivated in Agrawal et al. (2010).

The input *u* and resulting effect *y*^0^ are shown in Fig. 4 (black) together with the noise-corrupted observation *y* (gray).

## 4. LOCAL IDENTIFIABILITY

### 4.1. Practical identifiability at the true parameter

Unless pole-zero cancellations occurs (as further explained in Sec. 5.2) the system (1)–(2) is structurally identifiable from *u, z*. Since *h* (*z*) of (5) is strictly monotonous in *z* it also holds that the dynamics are structurally identifiable from *u, y*^0^.

However, the input *u* of Fig. 4 comprises merely two step changes. Furthermore, the experiment duration is less than one sixth of the slowest pole time constant of the true model (cf. table 1).

Increasing input excitation or prolonging the experiment would both be associated with ethical concerns related to patient safety. It is therefore natural to ask whether the parameters of (1)–(5) are *practically* identifiable from *u, y*.

Let therefore *y* = [*y*_1_ … *y*_*n*_]^T^ be a vector of the simulated observations, with *u* =[*u*_1_ … *u*_*n*_]^T^ being the corresponding infusion rates. Since *u* remains constant between consecutive samples, we can express the observation model as

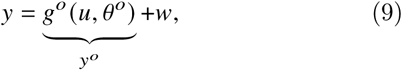

where *g*^*o*^ denotes the *true* model structure dynamics of (1)–(5), *θ* is a vector comprising of the parameters of table 1 (in that order), and *θ*^*o*^ signifies the true parameter values. Given the known initial condition *x*(0) = 0, *g*^*o*^ can be evaluated approximation-free at *u, θ*^*o*^ through zero-order-hold (ZOH) sampling.

The data distribution is defined by the probability density function (PDF) *p(y*|*θ)* and denotes the likelihood of observing *y* given *θ*. Therefore, the data distribution is also referred to as the likelihood of *0, L (θ)* = *p*(*y*|*θ*). Note that both *g* and *θ* of (9) are assumed to be deterministic, and so the likelihood of *θ* given the data, or observation, *y* is defined by the PDF of *w*.

Since *w* is IID with *w* ∼ 𝒩 (0, *σ*^2^), we have that

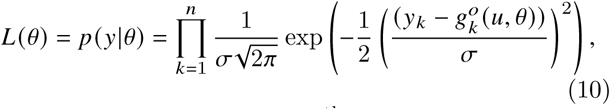

where 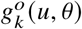 denotes the *k*^th^ element of the vector *g*^*o*^ *(u, θ)*. Maximizing the likelihood *L (θ)* of (10) is equivalent to maximizing the log-likelihood

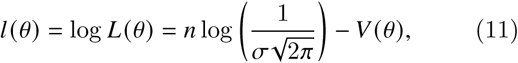

where the loss (or cost) function *V* is given by

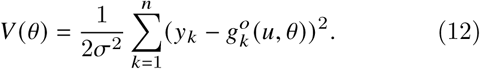

If 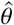 is an unbiased estimator of *θ*, meaning that 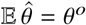, then the Cramér Rao Lower Bound (CRLB) expresses a lower bound on its variance:

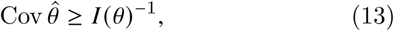

where the Fisher Information Matrix (FIM) is given by

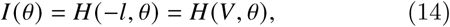

and where the Hessian *H* of a function *f* with respect to an argument *x* is a matrix with elements

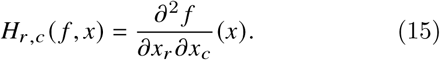

For the maximizer *θ = θ*^*o*^ of *l(θ)* the CRLB is tight. See e.g. Kay (1993) for derivation of the CRLB and discussion of necessary, and here fulfilled, regularity conditions for it to hold.

We can note that *V* of (12) conveys information about how well the model output *ŷ* (*u, θ*) = *g*^0^(*u,θ*) resembles the observation *y*. Particularly, the root mean square (RMS) error is 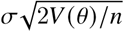.

The Hessian (14) also appears in the Taylor series expansion

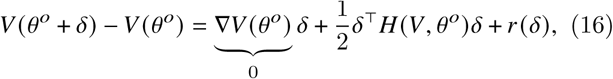

where the true parameter *θ*^*o*^ has been perturbed by *δ*, and where the residual *r* (*δ*) is a linear combination of monomials in the components of *δ*, each with degree of at least 3. If ‖ *δ* ‖ _2_ is small, the contribution of *r(δ)* to *V* is small.

The function *g*^*o*^(·, *θ*) has continuous second derivatives with respect to components of *θ*, resulting in *H* being symmetric with singular value decomposition (SVD) 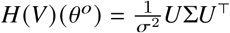. The CRLB states that the principal parameter space direction variance at the true parameter is lower bounded by the diagonal elements of *σ*^2^ Σ^−1^, since *I (θ*^*o*^) = *σ*^2^ (*U*Σ*U*^T^) ^−1^ = *σ*^2^*U* Σ^−1^*U*^T^. (If any diagonal element of Σ is zero, the estimate variance in the principal parameter space direction defined by the corresponding column of *U* will thus be infinite.)

For convenience of interpreting the sensitivities, we have normalized the parameter vector by the true parameter values, so that the true normalized parameter vector corresponding to (17)–(18) is 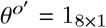, where ′ signifies the normalization and 1_8 × 1_ is a row vector of eight ones. Upon this normalization, the SVD matrices at the true parameter values become

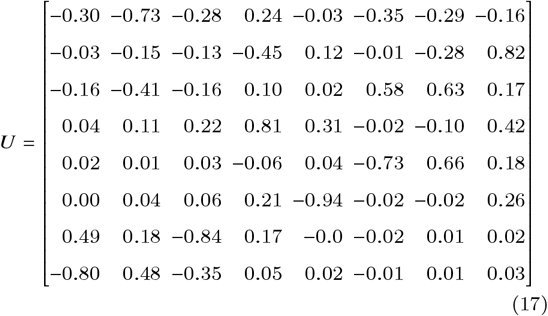

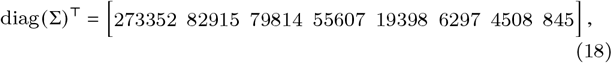

where the parameter order is that of table 1. The condition number of the FIM, being the ratio between the largest and smallest element in *σ*^2^Σ^−1^ is large, cond 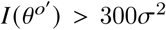, indicating high noise sensitivity and the last column of *U* in (17) reveals that the least certain parameter space direction at 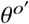 is roughly 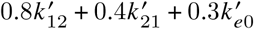.

## 5. GLOBAL IDENTIFIABILITY

### 5.1. Flexibility of parameterization

The analysis of Sec. 4.1 characterizes *local* identifiability of linear combinations of the parameters at *θ*^*o*^.

To analyze global identifiability of the individual parameters, we fixed them—one at a time—to a value that was off by a factor of 100 to investigate how well the remaining parameters could compensate for this. The quadratic loss *V* of (12) was minimized using Newton’s method with line search. Automatic differentiation was used to obtain approximation-free gradient and Hessian information, see Revels et al. (2016). After careful initialization, the model fits of Fig. 2 were obtained. The legend notation Δ: indicates which parameter has set to be off by a factor 100, and Δ*k*_13_ corresponds to a reduced-order model, detailed in Sec. 5.2. Corresponding Bode diagrams are shown in Fig. 3.

**Fig. 1.**
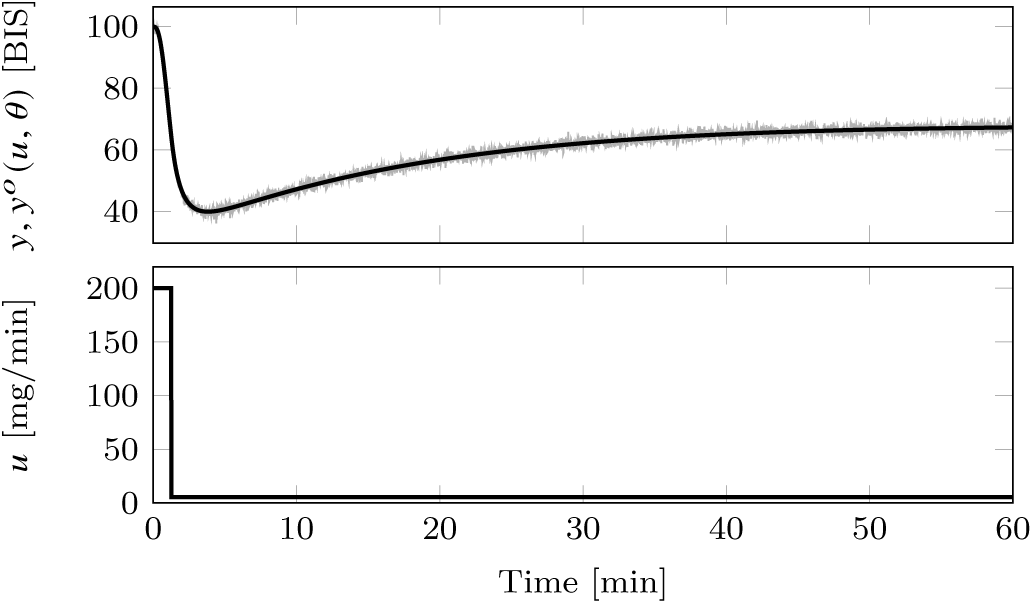
*Top:* Simulated true effect *y*^0^ = *g*^*o*^(*u, θ*^*o*^) (black) and noise-corrupted observation *y* (gray). *Bottom:* Propofol infusion profile *u*.

**Fig. 2.**
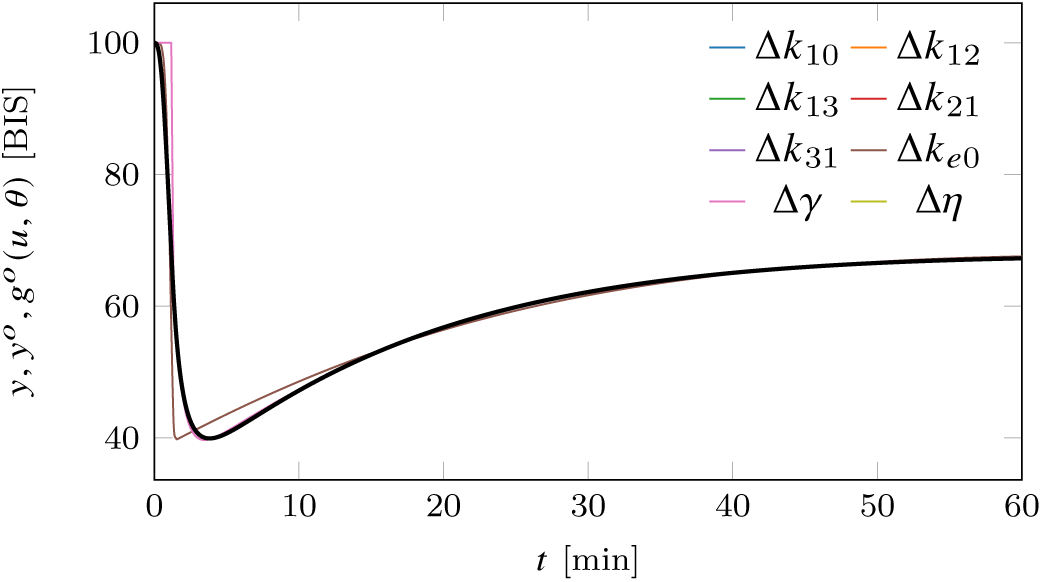
Simulated true effect *y*^0^ (black) where remaining curves show model outputs *g*^*o*^(*u,θ*) where all parameters (elements of *θ*) have been optimized one at a time (see legend). *k*_10_, *k*_21_, *k*_31_, *k*_*e*0_ and *γ* was fixed to a factor 100 times its true value and *k*_12_, *k*_13_ and *η* to 1/100 of its true value.

**Fig. 3.**
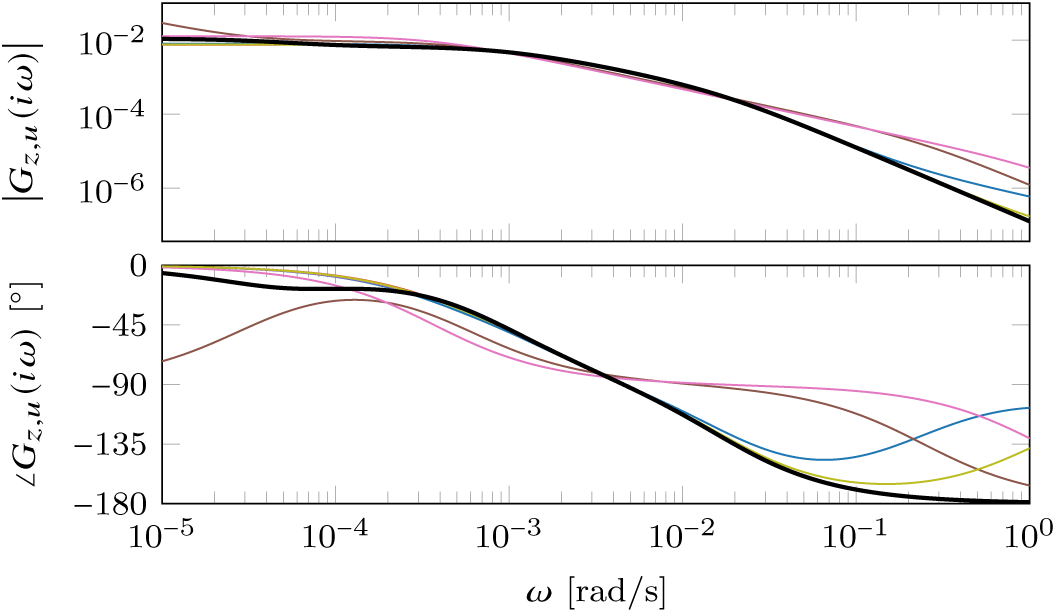
Bode plots of the models in Fig. 2. See Fig. 2 for further specification of the individual models.

Considering a model *g*(*u, θ*) where *g* ≠ *g*^*o*^ or *θ* ≠ *θ*^*o*^ leads to a bias error *y*_0_(*θ):*

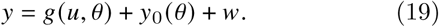

This results in a typically non-Gaussian residual *ϵ* = *y*_0_ *(θ)* +*w*, and consequently minimizing the loss *V*, corresponding to minimizing *ϵ*^T^*ϵ*, does in general no longer yield a *θ* that maximizes the log-likelihood *l* of the data. However, as seen in Fig. 2, the bias for all cases but Δ_*ke*0_ is negligible. Consequently, the relation (11) between *V* and *l* holds with good approximation for these cases. For Δ*k*_*e*0_, and to a lesser extent Δ*γ*, the fitting errors in Fig. 2 can be visually detected. This corresponds to some degree of identifiability, but reducing the error factor to 8 for *k*_*e*0_ and 2 for *γ* enables fitting models that are visually indistinguishable from the true dynamics in Fig. 2.

Inspired by the reduced-order models proposed in da Silva et al. (2012) we also minimized the quadratic loss *V* of (12) for a reduced-order model with three asymptotically stable real poles and one asymptotically stable zero. It is the result of this minimization plotted as Δ*k*_13_ in Fig. 2, and it naturally raises the question of which reduced-order models correspond to (3) through pole-zero cancellation, and what flexibility of the parameterization such cancellation provides.

### 5.2. Pole-zero cancellations

Here we characterize conditions imposed on the poles and zeros of reduced-order transfer functions, for them to constitute a realization of (1)–(2) with *θ* ≻ 0 and what flexibility, i.e. loss of identifiability, the cancelled dynamics provide.

It follows directly from the compartmental structure of (1)–(2) that *θ* ≻ 0 implies that *G*_*z,u*_*(s)* has strictly positive static gain, and that all its poles and zeros are real negative numbers (asymptotically stable). Before commencing the analysis we also note that there is a symmetry in that indices 2 and 3 of (1) can be interchanged.

*No cancelled dynamics* Assuming that no two of *k*_*e*0_, *k*_21_, *k*_31_ are equal, the static gain, zeros, and effect site pole are uniquely determined by *η, k*_21_, *k*_31_, and *k*_*e*0_, respectively. The characteristic equation (4c) then uniquely determines *k*_10_ = *p*_1_ *p*_2_ *p*_3_/(*k*_31_*k*_21_), and we can rewrite (4a)–(4b) as

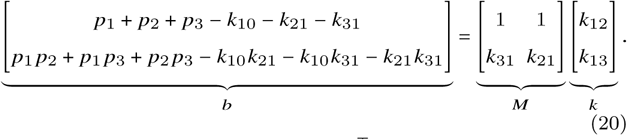

It is necessary that *b* = [*b*_1_ *b*_2_]^T^ ≻ 0 for all parameters to be positive. Since *k*_21_ ≠ *k*_31_, *M* has full rank and we are looking for a solution *k* = [*k*_12_ *k*_13_]^T^ = *M*^−1^*b* ≻ 0 of (20). We see from *b*_1_ = *k*_12_ + *k*_13_ that *k*_12_ < *b*_1_ (we only treat one of the two symmetric cases of *k*_12_ and *k*_13_ explicitly) is a necessary condition for the existence of such solution. With 0 < *k*_12_ < *b*_1_ we fulfill *b*_1_ = *k*_12_ + *k*_13_ with *k*_13_ = *b*_1_ − *k*_12_ > 0. We now rewrite the second characteristic equation as *b*_2_ = *k*_31_*k*_12_ + *k*_21_ (*b*_1_ − *k*_12)_ and solve for *k*_12_ =(*b*_2_ − *k*_21_*b*_1) /_ (*k*_31_ − *k*_21_). Due to symmetry we can exchange *k*_21_ with *k*_31_ to ensure *k*_12_ > 0. Combining the expression for *k*_12_ in known entities with the requirement *k*_12_ < *b*_1_ finally yields the necessary and sufficient condition

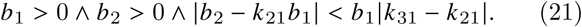

It is straightforward to expand the inequalities conditions into *p*_1_, *p*_2_, *p*_3_, *k*_10_, *k*_12_, *k*_13_, *k*_21_, *k*_23_, but this has been omitted here in the interest of space and readability.

#### 5.2.1. Single PK pole cancellation

Setting *k*_21_ = *k*_31_ results in the dynamics *x*_2_ and *x*_3_ of (1) becoming identical, and

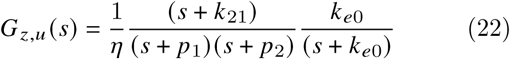

with characteristic equations *p*_1_ + *p*_2_ = *k*_10_ + *k*_12_ + *k*_13_ + *k*_21_ and *p*_1_ *p*_2_ = *k*_10_*k*_21_. The static gain, remaining zero and effect site pole are specified as in the cancellation-free case. This leaves the remaining parameters to determine the two remaining poles. As before, (4c) determines *k*_10_ = *p*_1_ *p*_2_ / *k*_21_. Inserting this expression, the first characteristic equation can be written *b* = *p*_1_ + *p*_2_ − *k*_21_ − *p*_1_ *p*_2_ / *k*_21_ = *k*_12_ + *k*_13_, and as before *b* > 0 constitutes a necessary condition. With it fulfilled and with *k*_12_ < *b*, 0 < *k*_13_ = *b − k*_12_ solves the first characteristic equation. The necessary and sufficient condition is thus 0 < *k*_12_ < *b* > 0. Relatedly, we see directly from (1) that cancelling two of the PK poles would require *k*_12_ = *k*_13_ = 0, which would void the strictly positive parameter requirement.

#### 5.2.2. Double cancellation

Setting *k*_21_ = *k*_31_ = *k*_*e*0_ results in two pole-zero cancellations in (3) and the transfer function

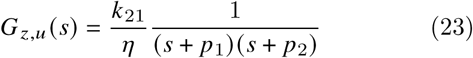

with characteristic equations identical to the PK compartment cancellation case. Solving the second characteristic equation for *k*_10_ and inserting into the first results in the necessary and sufficient realization condition *b* = *p*_1_ + *p*_2_ − *p*_1_ *p*_2/_ *k*_21_ − *k*_21_ > 0 ∧ *b* > *k*_12_ with *k*_13_ = *b* − *k*_12_ (or symmetric case by exchanging *k*_12_ for *k*_21_). The first inequality is quadratic in *k*_21_, and the corresponding quadratic equation has solutions *k*_21_ = *p*_1_ and *k*_21_ = *p*_2_. Since all parameters are positive, the inequality can thus be written min(*p*_1_, *p*_2_) < *k*_21_ < max(*p*_1_, *p*_2_). Solving *∂b/∂ k*_21_ = 0 we can also see that *b* (*k*_21_) is maximized at *k*_21_ = (*p*_1_ + *p*_2_)/2.

Again, it is straightforward to insert numerical values from the example to see within what bounds individual parameters could be varied without affecting *G* _*z,u*_.

## 6. DISCUSSION

It should not come as a surprise that the eight individual parameters of *θ* cannot be reliably identified from an experiment with poor input excitation (essentially the sum of an impulse and a step) that is furthermore short compared to the slowest pole dynamic. Particularly, loss of identifiability of the static gain is expected from an experiment duration less than one eighth of the slowest pole time constant of the true system (see Fig. 4 and table 1). A clear illustration of this is provided in Fig. 4, that shows the continuations the experiments in Fig. 2 under constant *u*.

**Fig. 4.**
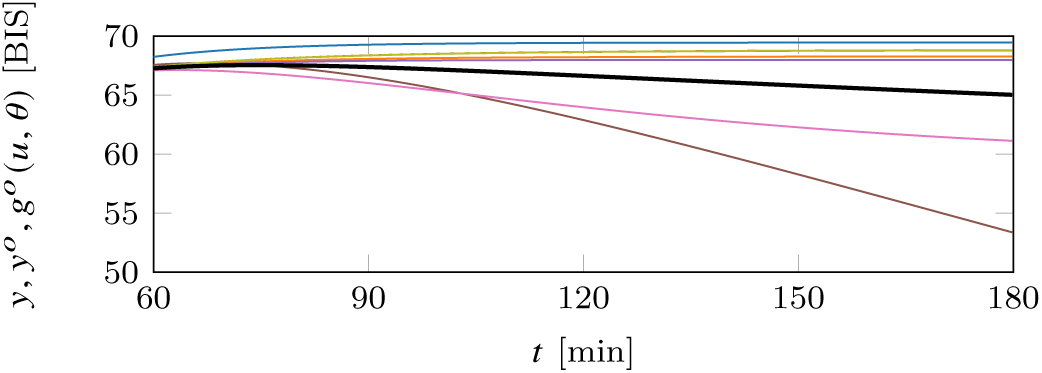
Continuation of the simulations in Fig. 2 beyond the first hour used to fit the models. See Fig. 2 for further specification of the individual models.

Yet, several factors worsen the prospect of identifiability from *real* data. Firstly, we have here assumed that the data was generated by dynamics that are in the set of the considered model class and that no disturbances act on the system in addition to a Gaussian IID measurement noise of known variance. Looking at EEG responses from actual patients, available through for example Eleveld et al. (2018) one quickly realizes that each of these assumptions is far from realistic. Co-administration of drugs also influences the anesthetic depth. Notably, opioids (administered for their analgesic effect) typically decrease *y*, see Vuyk (1997). The extent by which nociceptive stimuli affect *y* can to some extent be controlled using analgesic drugs, but not to an extent that their influence can be neglected. Secondly, the observation model (7) is a simplification mainly in that EEG monitors introduce considerable time lag caused by internal signal processing. For the NeuroSense, the lag dynamics are LTI and constituted by ZOH sampling of a second order filter with real pole time constants of 8 s, at *T*_*s*_ = 1 s, as further explained in Bibian et al. (2011). For the much more commonly applied BIS monitor (Medtronic, Dublin, Ireland), the corresponding filtering dynamics are proprietary, and experiments reported in e.g. Lee et al. (2019); Bibian et al. (2011) have demonstrated that they exhibit nonlinear rather than LTI behavior.

The two principal alternatives available to improve the situation is to use simplified models, as proposed in for example da Silva et al. (2012), Wahlquist et al. (2020), van Heusden et al. (2013), and account for the associated bias by designing controllers with a higher degree of robustness to model uncertainty to account for this bias.

The described situation arises in many domains when complex dynamics are to be identified from non-informative data. In our case, identification of the full PKPD structure results in low bias (at least for simulated data) but high variance (uncertainty) of the parameter estimates. Reduced-orders models can reduce this variance, but at the cost of increased bias.

This brings up the central question of what the model will be used for. Historically the closed-loop anesthesia control research community has relied on the PKPD structure, perhaps mainly because models reported in the literature have that structure. However, if the purpose of the model is feedback controller design, there is no apparent benefit in sticking to the mechanically motivated model structure of (1)–(2). Therefore, it can be viewed as good news that comprehensive original experimental datasets for propofol PK and PD modeling have lately been compiled, perhaps most notably in the supplement of Eleveld et al. (2018). Setting out in such data it is for the first time possible to combine the activities of modelling and controller design, thus eliminating the reason to guess (or neglect) what uncertainties to associate with published PKPD models when considering them as a basis for controller design.

## 7. CONCLUSION

The classic PKPD model structure for propofol is not practically identifiable from induction phase data available in the clinic. Reliable individualization of drug delivery, beyond what is possible through compensating for demographic trends that can be determined *a priori*, will therefore need to be achieved by other means. The recent availability of comprehensive original propofol PKPD modeling datasets opens up for the identification of other model structures. If the objective is closed-loop control, it is more important for the dynamical patient model to be accurate around the intended closed-loop bandwidth than for it to maintain a physiologically motivated structure. Particularly, a closer investigation on how control performance is affected by employing less flexible models (with fewer parameters), is motivated in the light of the investigated identifiability issues.

## 8. ACKNOWLEDGEMENTS

The authors would like to acknowledge Johan Grönqvist, Lund University, for help with the implementation of the pharmacological model. This work was partially funded by the Swedish Research Council (grant 2017-04989). The authors are members of the ELLIIT Strategic Research Area at Lund University.

## Footnotes

1 The resolution in *t* is typically ≈ 1 s and the resolution in *u* for a standard 10 mg/ml propofol solution is ≈ 0.02 mg/min, to be compared with the input magnitude and time scales of a representative therapeutic profile in Fig. 4.

2 In the literature, the input matrix of (1) is often multiplied by *EC*_50_, while (5) and (6) are combined into the standard Hill function form 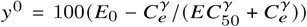. The input-output behavior becomes equivalent, but the normalization of the internal signals *z* and *h* (*z*) is lost.

3 The aforementioned discrepancy between system time constants and sampling period allows for lowpass filtering before subjecting *y* to feedback control, making *σ*^2^ = 9^2^ BIS a somewhat pessimistic assumption in that context.

4 The infusion rate corresponds to 1200 ml/h (being the max rate of e.g. the Alaris TIVA infusion pump, (BD, Franklin Lakes, NJ)) with 10 mg/ml propofol (being a common emulsion concentration).

